# Active avoidance requires inhibitory signaling in the rodent prelimbic prefrontal cortex

**DOI:** 10.1101/241943

**Authors:** Maria M. Diehl, Christian Bravo-Rivera, Jose Rodríguez-Romaguera, Pablo A. Pagán-Rivera, Anthony Burgos-Robles, Gregory J. Quirk

**Author notes:** Corresponding Author: Maria M. Diehl, Department of Psychiatry and Neurobiology & Anatomy, University of Puerto Rico School of Medicine, San Juan, PR 00936. **Competing interests**: The authors declare that no competing interests exist.

## Abstract

Much is known about the neural circuits of conditioned fear and its relevance to understanding anxiety disorders, but less is known about other anxiety-related behaviors such as active avoidance. Using a tone-signaled, platform-mediated active avoidance task, we observed that pharmacological inactivation of the prelimbic prefrontal cortex (PL) delayed initiation of avoidance. However, optogenetic silencing of PL neurons did not delay avoidance. Consistent with this finding, inhibitory, but not excitatory, responses of rostral PL neurons to the tone were correlated with initiation of avoidance. To oppose inhibitory responses, we photoactivated rostral PL neurons during the tone to maintain pre-tone firing rate. Photoactivation of rostral PL (but not caudal PL) neurons at 4 Hz (but not 2 Hz) delayed or prevented avoidance. These findings suggest that the initiation of active avoidance requires inhibitory neuronal responses in rostral PL, and underscores the importance of designing behavioral optogenetic studies based on neuronal firing patterns.

## Introduction

Core symptoms of post-traumatic stress disorder and other anxiety disorders include excessive fear and maladaptive avoidance (DSM-V, 2013). The neural mechanisms of excessive fear have been well-characterized in rodents using Pavlovian fear conditioning (Johansen et al., 2011, Herry and Johansen, 2014, Giustino and Maren, 2015, Do Monte et al., 2016), yet the neural underpinnings of active avoidance are just beginning to emerge. Previous work in rats has shown that the prefrontal cortex, amygdala, and striatum are all necessary for the expression of active avoidance (Martinez et al., 2013, Moscarello and LeDoux, 2013, Beck et al., 2014, Jiao et al., 2015, LeDoux et al., 2017). Using a tone-signaled, platform-mediated avoidance task, we previously observed that pharmacological inactivation of the prelimbic prefrontal cortex (PL) impaired the expression of avoidance without affecting freezing (Bravo-Rivera et al., 2014). Furthermore, avoidance that persisted following extinction training was correlated with excessive PL activity, as indicated by the immediate early gene cFos (Bravo-Rivera et al., 2015), suggesting that PL activity drives active avoidance.

Important questions remain, however, regarding the role of PL in avoidance. First, how do PL neurons signal avoidance? Fear conditioning mainly induces excitatory responses to conditioned tones in PL that correlate with freezing (Baeg et al., 2001, Burgos-Robles et al., 2009, Sotres-Bayon et al., 2012, Isogawa et al., 2013, Pendyam et al., 2013, Chang et al., 2010), but firing properties of PL neurons during active avoidance have not been previously studied. In platform-mediated avoidance, PL signaling of avoidance may differ from PL signaling of freezing or foraging for food (Burgos-Robles et al., 2013), both of which can interfere with platform avoidance. Second, is PL activity correlated with the initial decision to avoid and/or the subsequent expression of avoidance?

We addressed these questions with single unit recordings to determine PL signaling during discrete events leading to the successful execution of active avoidance. We then optogenetically silenced or activated PL neurons, based on the observed firing properties. We find that inhibitory responses (rather than excitatory responses) of rostral PL neurons at tone onset are correlated with the initiation of active avoidance. Reducing these inhibitory neuronal responses with photoactivation delayed or prevented the initiation of avoidance, suggesting that prefrontal inhibition underlies the decision to avoid danger.

## Results

### Pharmacological inactivation of PL delays the initiation of avoidance

We first replicated prior findings that pharmacological inactivation of PL with muscimol (MUS) impairs avoidance in this task (Bravo-Rivera et al., 2014), with two modifications: 1) we used fluorescently labeled MUS to assess spread to adjacent regions, and 2) we analyzed the time course of avoidance behavior across the 30 sec tone cue. Because the 2 sec shock co-terminates with the tone, the rat has 28 sec to stop pressing the lever for food and step onto the platform to escape the shock. Furthermore, in this task, avoidance comes at a cost, as it competes with access to food. Thus, the involvement of PL could vary with changes in the cost and/or urgency of avoidance as the tone progresses (Zeeb et al., 2015, Hosking et al., 2016).

Histological analysis showed that MUS was confined to PL in its mid rostral-caudal extent (Figure 1A). Rats showing substantial spread to adjacent infralimbic cortex were excluded (n=3). In some cases, MUS reached the ventral half of cingulate cortex (Cg1), but they were included due to similar functions of Cg1 and PL (Courtin et al., 2014). Following surgical implantation of cannulas, rats were trained in platform-mediated avoidance over 10 days as previously described (Figure 1B, Bravo-Rivera et al., 2014, Rodriguez-Romaguera et al., 2016). At Test 1 (Day 11), we infused MUS into PL at the same concentration as our prior studies using fluorescent MUS (Do-Monte et al., 2015b, Rodriguez-Romaguera et al., 2016) and waited 45 minutes before commencing a 2-tone test of avoidance expression (without shock). Figure 1C shows that MUS inactivation significantly reduced the time spent on the platform during the tone, as compared to saline (SAL) infused controls (SAL 92% vs. MUS 57%, unpaired t-test, t_(28)_=-4.019, p<0.01, Bonferroni corrected). An analysis of avoidance across the tone in 3 sec bins (Figure 1D) indicated that 11/13 MUS-infused rats were significantly delayed in their initiation of avoidance (repeated measures ANOVA, F_(1,9)_=4.076, p<0.001; post hoc Tukey test, 0–15 sec **p<0.01, 15–21 sec *p<0.05), and 2/13 rats never avoided (Mann Whitney U Test, p<0.001, Figure 1E). MUS also increased freezing during the tone (Figure 1E inset; SAL = 36% vs. MUS = 55% freezing, unpaired t-test, t_(28)_=2.460, p=0.020). MUS inactivation of PL had no effect on locomotion, as indicated by distance traveled in an open field test during a 5 min period (SAL n=10, 13.23 m vs. MUS n=10, 12.53 m, unpaired t-test, p=0.614). Nor did it affect anxiety levels, as both groups spent a similar amount of time in the center of the open field (SAL = 15.69 sec vs. MUS = 18.76 sec, unpaired t-test, p=0.363). Thus, in the majority of rats, pharmacological inactivation of PL slowed the initiation of avoidance during the tone.

**Figure 1.**
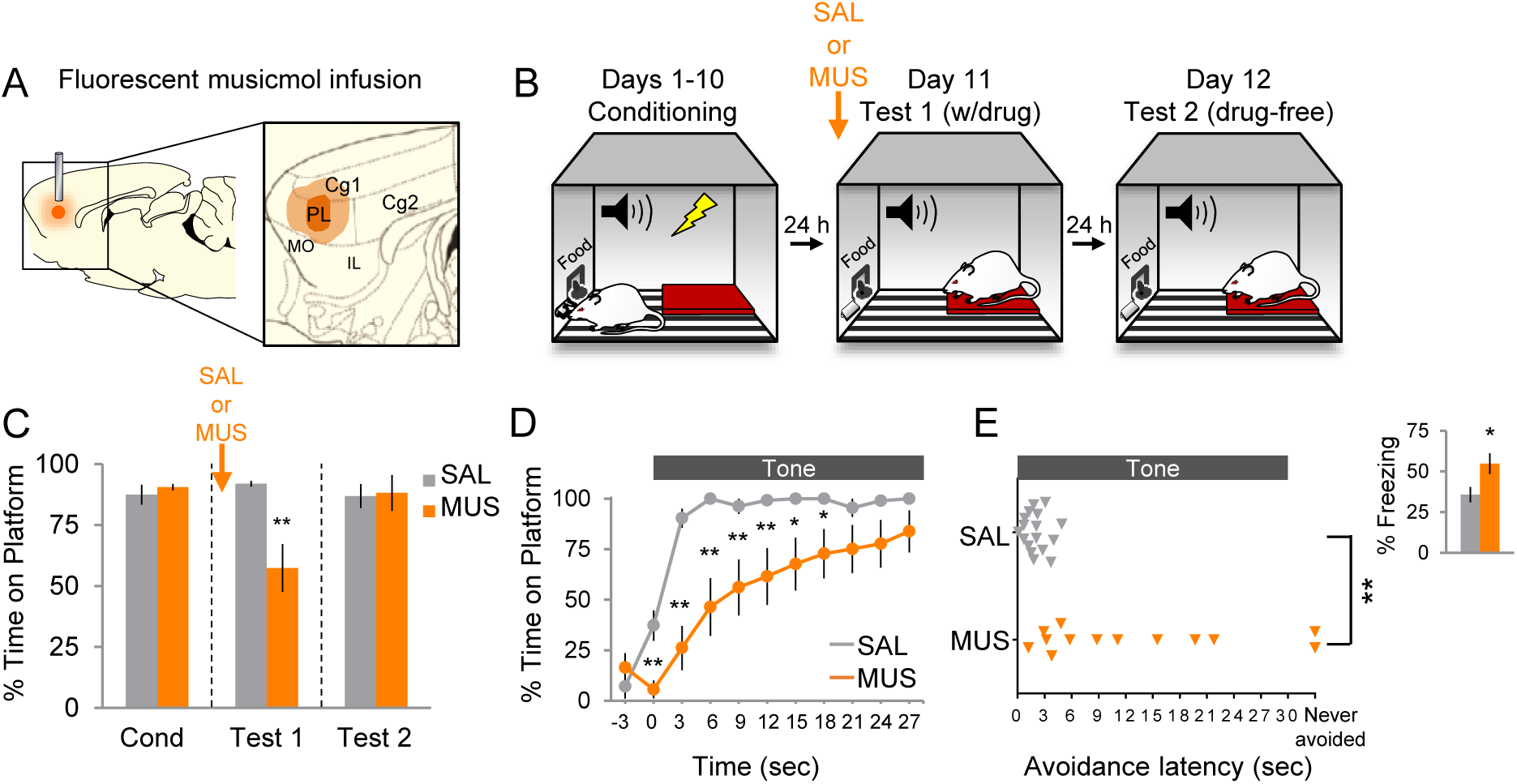
Pharmacological inactivation of prelimbic cortex delays avoidance. **A**. Schematic of MUS infusion showing min (dark orange) and max (light orange) extent of infusion in PL. **B**. Rats were trained across 10 days to avoid a tone-signaled foot-shock by stepping onto a platform. On Day 11, rats received 2 tone presentations (without shock) 45 min after MUS infusion. On Day 12, rats received a second 2-tone test drug free. **C**. Percent time on platform during Tone 1 on Days 10, 11 (with MUS), and 12 for saline controls (SAL, n=17; grey) and MUS rats (n=13, orange). **D**. Time spent on platform in 3 sec bins (Tone 1, Test 1) revealed that MUS rats were significantly delayed in their avoidance compared to SAL controls (repeated measures ANOVA, post hoc Tukey test). **E**. Latency of avoidance for each rat (Mann Whitney U Test, Tone 1, Test 1). *Inset*: Effect of MUS inactivation (Tone 1, test 1) on freezing during the tone (unpaired t-test). Data are shown as mean ± SEM; *p<0.05, **p<0.01.

### Photosilencing of PL neurons does not delay avoidance

Because pharmacological inactivation of PL delayed avoidance initiation, we reasoned that tone-induced activity in PL would be essential for early avoidance. To asses this, we used an optogenetic approach, expressing the microbial opsin archaerhodopsin (Arch) in PL, which causes a hydrogen proton efflux to hyperpolarize neurons when exposed to 532 nm (green) light (Chow et al., 2010, Han et al., 2011). We delivered Arch by infusing an adeno-associated virus (AAV) encoding both Arch and enhanced yellow fluorescent protein (eYFP) under the control of the CAMkII-α promotor to target glutamatergic projection neurons (AAV5:CaMKIIα::eArchT3.0-eYFP; Liu and Jones, 1996). We first confirmed that Arch silences PL neurons in anesthetized rats by recording extracellular activity in PL while illuminating Arch-expressing PL neurons (Figure 2A). Laser illumination significantly decreased the firing rate of 41/70 neurons, and increased the firing rate of 11/70 neurons (Figure 2A, paired t-tests comparing pre-laser vs laser activity of each unit using 1-sec time bins, all p’s<0.05).

**Figure 2.**
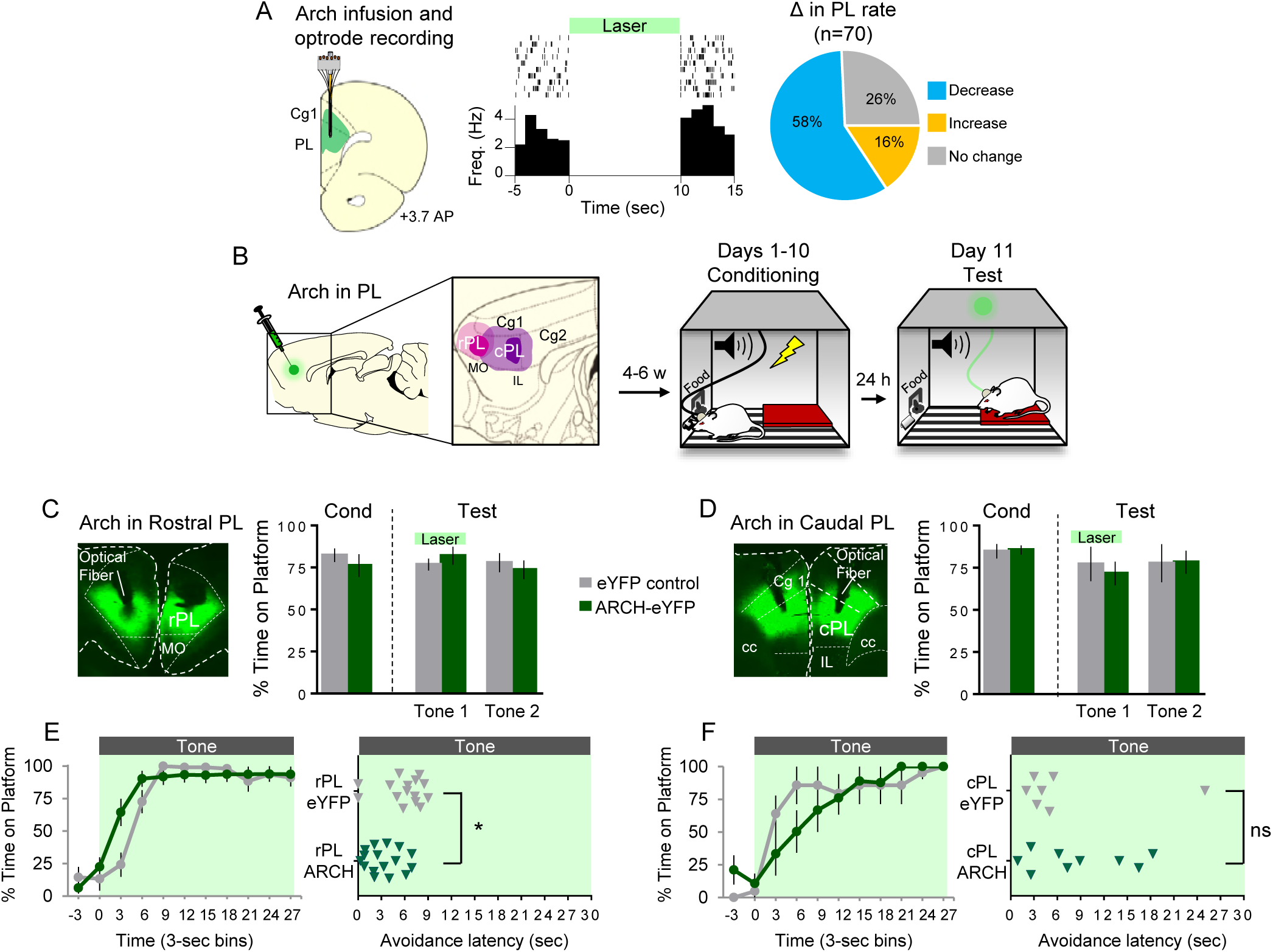
Optogenetic silencing of prelimbic neurons does not delay avoidance. **A**. *Left:* Schematic of Arch expression and optrode placement (n=2 rats). *Middle:* Rasters and peristimulus time histogram of a single PL neuron showing a decrease in firing rate during laser illumination (8–10mW, 532nm, 10s ON, 10s OFF, 10 trials). *Right:* Proportion of PL neurons that exhibited a decrease (blue, n=41), increase (gold, n=11), or no change (grey, n=18) in firing rate. **B**. Schematic of virus infusion, location of min/max expression of AAV in rPL (pink) and cPL (purple), followed by avoidance training and test. At Test, 532nm light was delivered to rPL or cPL during the entire 30-second tone presentation (Tone 1). **C**. *Left:* Micrograph of Arch expression and optical fiber placement in rPL. *Right:* Percent time on platform at Cond (Day 10, Tone 1) and Test (Day 11, Tone 1 with laser ON and Tone 2 with laser OFF) for eYFP-rPL control (n=15, grey) and Arch-rPL rats (n=17, green). **D**. *Left:* Micrograph of Arch expression and optical fiber placement in cPL. *Right:* Percent time on platform during Cond and Test for eYFP-cPL control (n=7, grey) and Arch-cPL rats (n=9, green). **E**. *Left:* Time spent on platform in3 sec bins (Tone 1 at Test) revealed no effect of silencing rPL-Arch neurons compared to eYFP controls (repeated measures ANOVA, post hoc Tukey test). *Right*: Latency of avoidance for each rat (Tone 1 at Test). rPL-Arch rats showed a decrease in avoidance latency (Mann Whitney U test, p<0.05). **F**. Timeline of avoidance (left) and latency (right) for cPL-eYFP control rats and cPL-Arch rats. All data are shown as mean ± SEM; p<0.05.

Next, we infused Arch into PL, distinguishing between rostral PL (rPL; defined as dorsal to medial orbitofrontal cortex and anterior to the infralimbic cortex) and caudal PL (cPL; defined as dorsal to the infralimbic cortex; Figure 2B) based on distinct connectivity of these sub-regions (Floyd et al., 2000, Floyd et al., 2001). Ten days of avoidance training commenced 6–8 weeks after viral infusion. Rats were then tested for avoidance expression. PL neurons were illuminated with laser during the first tone, followed by a second tone with the laser off. Surprisingly, avoidance initiation was not impaired by photosilencing of either rPL (Figure 2C) or cPL (Figure 2D&F). Further examination showed that photosilencing rPL neurons had no significant effect on the time course of avoidance following tone onset (Figure 2E left). However, rPL-Arch rats avoided significantly earlier compared to rPL-eYFP control rats, as measured by avoidance latency (Figure 2E right, Mann Whitney U test, p=0.0202).

The lack of impairment of avoidance may suggest that we failed to sufficiently inhibit PL activity via Arch photosilencing. Arguing against this, however, photosilencing rPL neurons during early avoidance training (on day 2) significantly reduced tone-induced freezing (eYFP-control, n=9, 31% vs. eYFP-Arch n=8, 7% freezing, unpaired t-test, t_(15)_=0.288, p=0.0115). Thus, contrary to our initial hypothesis, excitatory activity of PL projection neurons does not appear to be necessary for initiation of avoidance. Instead, silencing rPL tended to facilitate avoidance (as indicated by the decrease in latency), raising the possibility that avoidance signaling may involve rPL inhibition rather than excitation.

### Inhibitory responses in rostral PL neurons correlate with the initiation of avoidance

An assumption of our photosilencing approach was that increased activity in PL neurons is correlated with avoidance, however, this hypothesis had never been tested. We therefore performed extracellular single unit recordings in PL of well-trained rats during avoidance expression. Units were recorded from the full rostral-caudal extent of PL (Figure 3A). We first characterized PL activity at tone onset (Figure 3B-D). Both excitatory and inhibitory responses were observed at tone onset (Figure 2B right). Figure 3C shows the proportions of neurons that were significantly responsive (at each 500 ms bin) throughout the tone. Out of a total of 241 neurons, 34 were excited (14%, Z > 2.58 (p<0.01) in the first 500 ms) and 25 were inhibited (10%, Z < −1.96 (p<0.05) in the first or second 500 ms bin) at tone onset, relative to 10 sec of pre-tone activity (Figure 3D left). This brief post-tone latency (<1 sec) was selected to ensure that the activity of PL neurons was limited to the tone, and not subsequent behavior, such as platform entry, which typically occurred ~5 sec after tone onset (see black dots above graph in Figure 3C).

**Figure 3.**
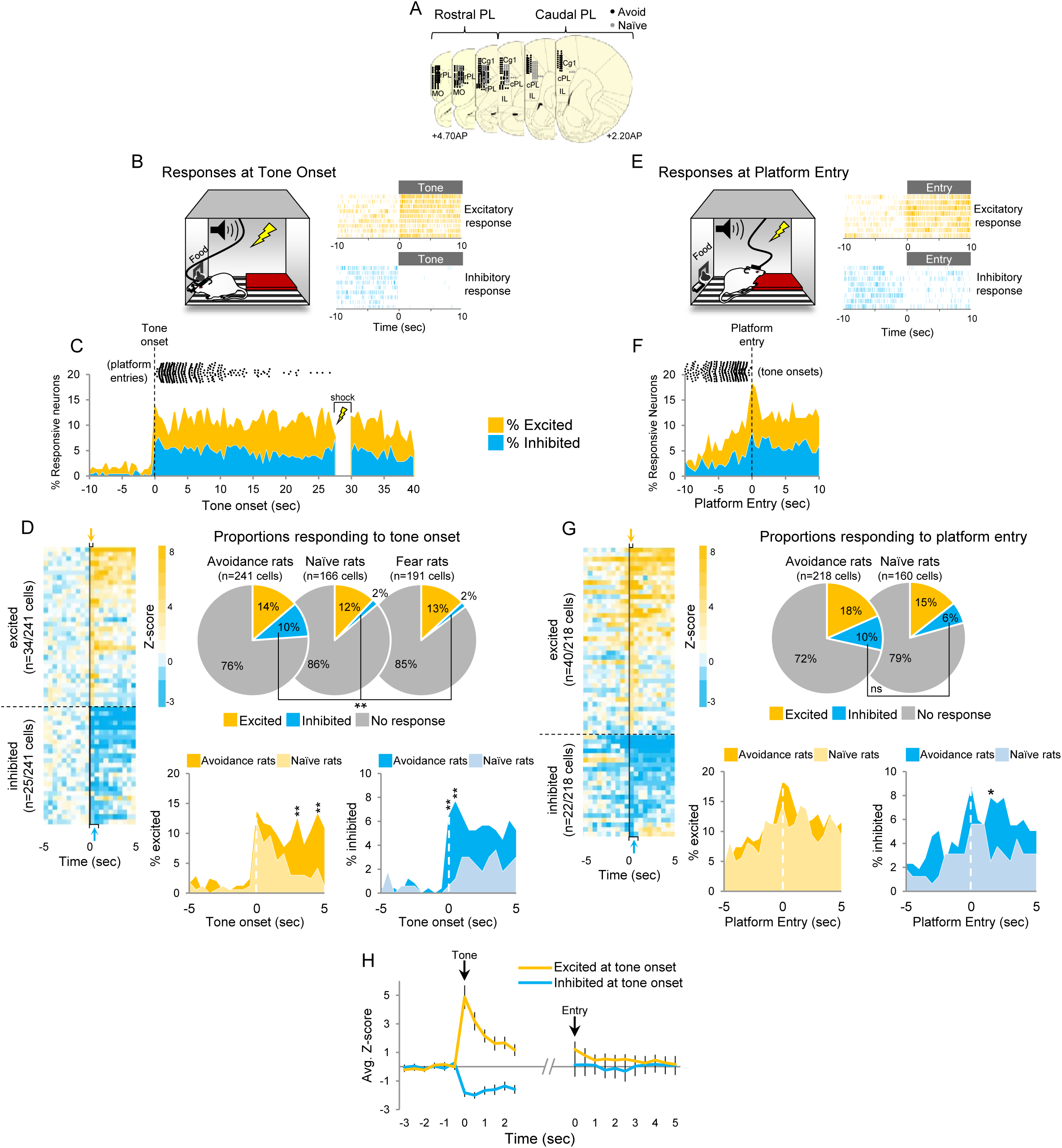
Initiation of avoidance is correlated with inhibition in rostral PL neurons. **A**. Location of recordings across PL (n=7 avoidance-trained and n=8 naïve rats). **B**. *Left:* Cartoon of rat behavior at tone onset during unit recordings. *Right:* single unit examples of an excitatory (gold rasters) and inhibitory tone response (blue rasters). Each row represents a single trial. **C**. Percentage of neurons that were excitatory (gold) or inhibitory (blue) throughout the tone. Time of platform entry (black dots), for all successful trials (n=340) in avoidance rats is indicated relative to tone onset. **D**. *Left:* Heat map of normalized responses (z-score) to tone onset (Time = 0 sec) of neurons in avoidance rats. Each row represents one neuron, bin = 0.5 sec. Arrows indicate bins used to determine criteria for excitatory (gold, first 500ms bin), or inhibitory (blue, first or second 500ms bin) tone responses. *Right:* Pie charts showing proportions of neurons that were excited, inhibited, or non-responsive at tone onset in avoidance (n = 34, 25, 182, respectively), naïve (n = 20, 3, 143, respectively), and fear conditioned rats (n = 25, 3, 163, respectively). Proportion of inhibitory responses were significantly greater in avoidance rats compared to naïve and fear rats (Chi Square test, **p<0.001). *Bottom:* Percentage of cells that were excited in avoidance (gold) or naïve (light gold) rats (left) or inhibited in avoidance (blue) or naïve (light blue) rats (right) around tone onset (Fisher exact tests). **E**. *Left:* Cartoon of rat entering platform after tone onset during unit recordings. *Right:* single unit examples of an excitatory (gold rasters) and inhibitory platform entry response (blue rasters). **F**. Percentage of neurons that were excitatory (gold) or inhibitory (blue) at platform entry. Time of tone onset (black dots), for all successful trials (n=340) in avoidance rats is indicated relative to platform entry. **G**. *Left*: Heat map of normalized responses to platform entry (Time = 0 sec) of neurons in avoidance rats. *Right:* Pie charts showing proportions of neurons that were excited, inhibited, or non-responsive at platform entry in avoidance (n = 40, 22, 156, respectively) and naïve rats (n = 23, 10, 127, respectively). *Bottom:* Percentage of cells that were excited in avoidance (gold) or naïve (light gold) rats (left) or inhibited in avoidance (blue) or naïve (light blue) rats (right) after platform entry (Fisher exact tests). **H**. Tone responsive neurons were not responsive to platform entry. All data are shown as mean ± SEM; *p<0.05, **p<0.01.

To determine if these tone responses were correlated with avoidance, rather than sensory perception of the tone, we compared PL responses in this group of rats with those of a naïve control group trained to press for food and presented with tones in the same chamber with the platform. Naïve rats were free to mount the platform and explore the chamber, but were never shocked. To determine whether activity at tone onset might represent the aversiveness of the tone, we also compared responses in avoidance rats with responses in rats subjected to auditory fear conditioning in the same chamber (re-analysis of data from Burgos-Robles et al., 2009). Surprisingly, there were no significant differences in the percentage of neurons showing excitatory tone responses in the avoidance group compared to the naïve or fear groups (Figure 3D right; avoidance-trained: 34/241 (14%), naïve: 20/166 (12%), fear: 25/191 (13%), Chi Square = 0.242, p=0.886). Inhibitory responses, however, occurred more frequently in avoidance-trained rats compared to naïve or fear rats (avoidance-trained: 25/241 (10%), naïve: 3/166 (2%), fear: 3/191 (2%), Chi Square = 22.649, p<0.001).

We then compared tone responses in avoidance and naïve rats at multiple time points around the tone onset. Consistent with the data shown in the pie charts of the avoidance and naïve groups, the percentage of excited cells did not significantly differ between avoidance (gold) and naïve (light yellow) rats at tone onset (time = 0 sec, Figure 3D bottom). The percentage of inhibited cells, however, was significantly higher in avoidance rats compared to naïve rats during the first two 500 ms bins (Fisher Exact tests, both p’s<0.01). Group differences observed after tone onset for both excitatory and inhibitory responses likely reflect the expression of avoidance behavior, which occurred in the avoidance group but not the naïve group.

We next examined PL activity at platform entry, defined as the moment at which the rat’s head entered the platform zone (Figure 3E-G). Activity around platform entry was compared to the same pre-tone baseline used in assessing responses to tone onset. Both excitatory and inhibitory responses to platform entry were observed (Figure 3E right). Figure 3F shows the proportion of neurons that were responsive at each 500 ms time bin around platform entry. PL neurons were either excited (n=40/218; 18%; Z > 2.58 in the first 500 ms) or inhibited (22/218; 10%; Z < −1.96 in the first or second 500 ms bin) at platform entry (Figure 3G left). The proportions showing each response in avoidance-trained rats did not differ significantly from those in naïve controls (Figure 3G right; n=23/160; 15% excited; n=10/160; 6% inhibited, Fisher Exact Tests, excited p=0.331; inhibited p=0.197), suggesting that platform entry responses in PL represent sensory perception and/or motor responses rather than avoidance of threat (Amir et al., 2015). Only inhibitory responses at tone onset correlated with initiation of avoidance of threat.

We then compared platform entry responses in avoidance and naïve rats at multiple time points around platform entry. Similar to tone responses, the percentage of excited cells did not significantly differ between avoidance (gold) and naïve (light yellow) rats at platform entry (time = 0 sec, Figure 3G bottom). Nor did the percentage of inhibited cells significantly differ at most time points (Fisher exact test), suggesting that activity changes at platform entry do not reflect avoidance of threat. Group differences observed after platform entry likely reflect sustained tone responses in avoidance rats, which were not present in naïve rats, as they mounted the platform outside of the tone.

In order to determine whether platform entry responses were distinct from tone responses, we examined activity during platform entry only in the neurons that showed tone responses (Figure 3H). Of the 34 cells that were excited at tone onset, only 7 (20%) were also excited at platform entry. Similarly, of the 25 cells that were inhibited at tone onset, only 9 (36%) were also inhibited at platform entry. Figure 3H shows the average normalized response (Z-score) of excited or inhibited responses at tone onset, followed by their responses at platform entry. This demonstrates that neurons exhibiting platform entry responses are largely distinct from those exhibiting tone onset responses. Taken together, these results show that initiation of avoidance is correlated with inhibitory responses in PL at tone onset but not with excitatory or inhibitory responses at platform entry.

Further characterization of inhibitory tone responses in PL revealed that most inhibitory responses (n=20/25) were not sustained 15 sec after tone onset, whereas 20% (n=5/25) were sustained throughout the tone (Figure 4A). Inhibition reduced the firing rate from an average of 6Hz to 2 Hz within 1 sec after tone onset (Figure 4B). The majority of neurons showing inhibitory tone responses were located in rPL (blue, n=22/25) rather than cPL (Figure 4C; purple; n=3/25, Fisher Exact Test, p=0.0383) and were likely putative projection neurons, based on their spike width and baseline firing rate (> 225 μs, <15 Hz for PL) as shown in Figure 4D (for method, see Sotres-Bayon et al., 2012). These results suggest that inhibition of rPL projection neurons at tone onset signals the initiation of active avoidance.

**Figure 4.**
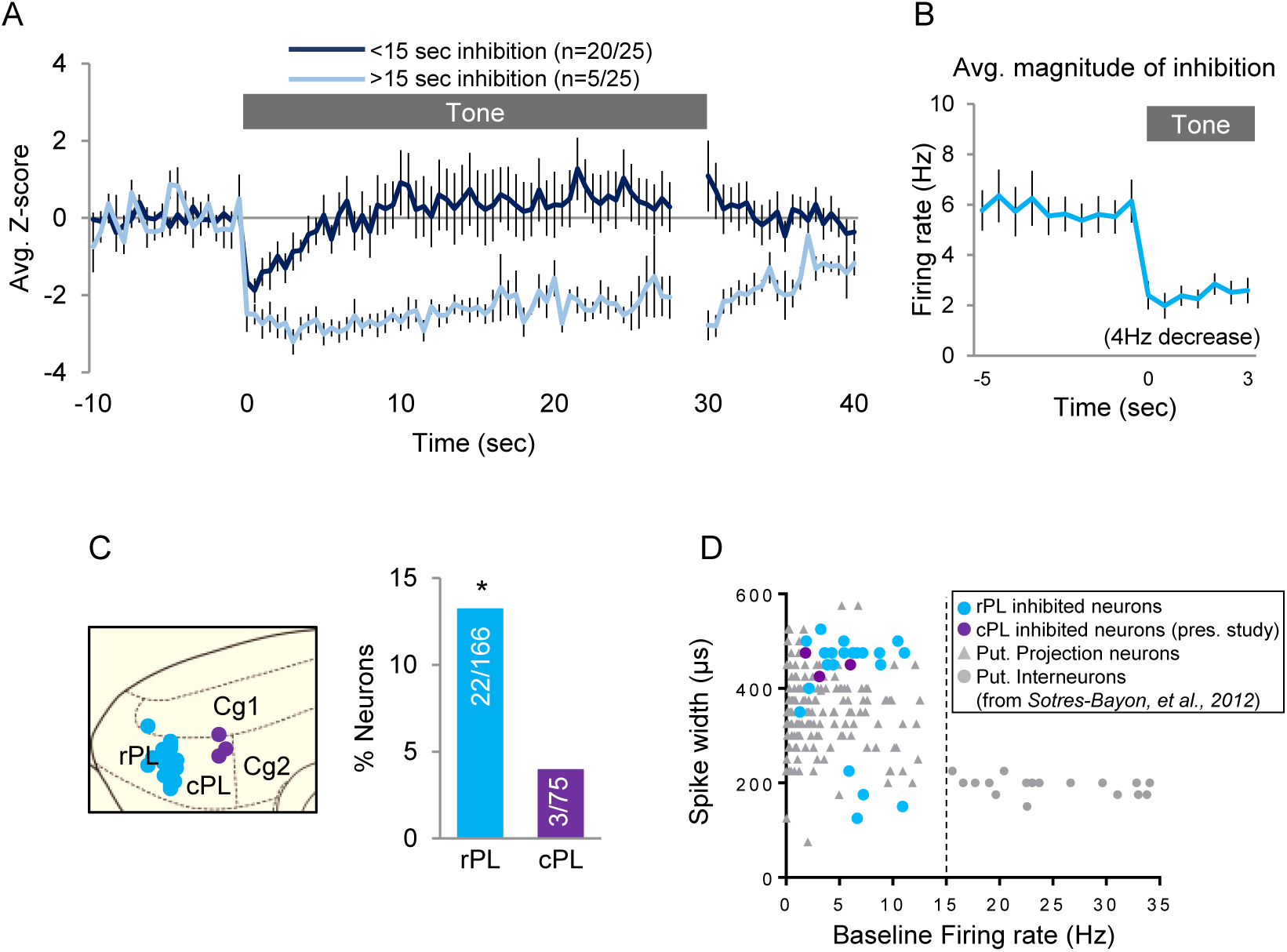
Characterization of inhibitory tone responsive neurons. **A**. Normalized firing rate of cells that were inhibited for less than 15 sec (dark blue) or more than 15 sec (light blue) after tone onset. **B**. Average inhibitory response of neurons decreased from a baseline firing rate of 5.8 Hz to 1.98 Hz at tone onset. **C**. *Left:* sagittal view of location of inhibitory tone responsive neurons in rPL (blue) or cPL (purple). *Right:* Histological analysis revealed that more inhibited neurons were located in rPL (Fisher Exact Test). **D**. Classification of PL neurons into putative projection neurons (gray triangle) or interneurons (gray circle) based on spike width and baseline firing rate (Sotres-Bayon et al, 2012). Neurons showing inhibitory responses (blue/purple, n=25) were likely projection neurons. All data are shown as mean ± SEM; *p<0.05.

### Countering inhibitory responses in rostral PL neurons delays or prevents avoidance

Our recording data demonstrate that inhibitory tone responses in rPL correlate with initiation of avoidance. Because inhibited neurons decreased their firing rate from 6 Hz to 2 Hz on average (Figure 4B), we reasoned that opposing this decrease should impair initiation of avoidance. To oppose inhibition, we used channelrhodopsin (ChR2) targeting CAMkIIα-positive neurons to activate rPL neurons at 4 Hz, concurrent with the tone. To confirm our method, we first measured extracellular unit activity in anesthetized rats from ChR2-expressing rPL neurons exposed to blue light (473nm) illumination (Figure 5A). Figure 5B shows a representative rPL neuron increasing its firing rate to 4 Hz photoactivation. We found that 4 Hz photoactivation increased the firing rate in 38% of the neurons and decreased the firing rate in 24% of neurons (Figure 4C left; n=112, 4 Hz, 30 sec duration, 5 ms pulse width, 8–10 mW illumination, student’s t-test, p<0.05). Photoactivation at 4 Hz increased firing rate from 1.41 Hz to 3.34 Hz on average, in neurons that were significantly excited (Figure 4C right). We found that photoactivation at 2Hz increased the firing rate in 46% and decreased the firing rate of 34% of neurons (Figure 4E left; n=76, 2 Hz, 30 sec duration, 5 ms pulse width, 8–10 mW illumination, student’s t-test, p<0.05). Photoactivation at 2 Hz increased firing rate from 0.4 Hz to 1.16 Hz on average, in neurons that were significantly excited (Figure 4E right). As expected, 2 Hz photoactivation had a weaker effect than 4Hz activation on driving spiking activity in rPL neurons.

**Figure 5.**
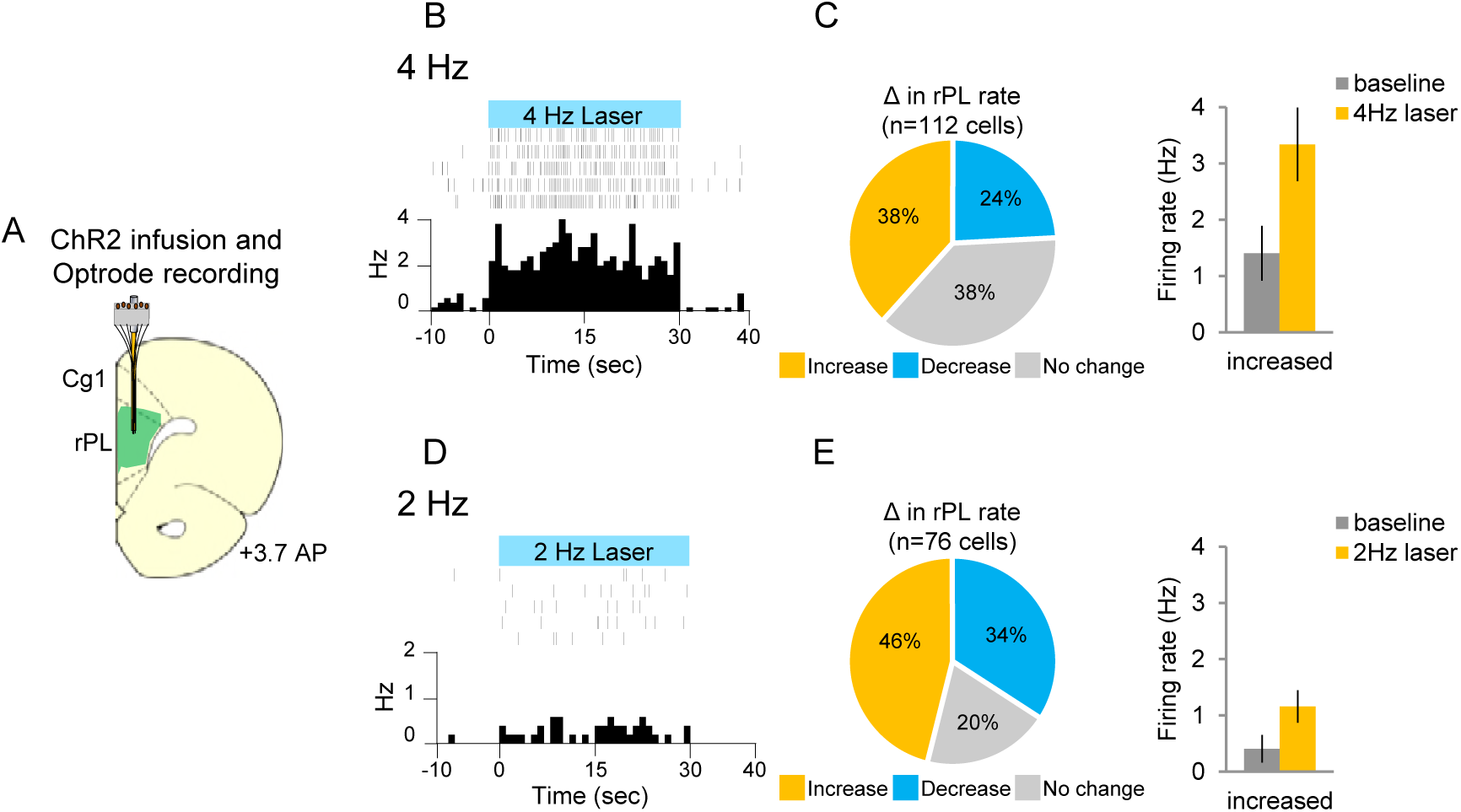
4 Hz photoactivation and single unit recordings of rostral PL neurons in anesthetized rats. **A**. Schematic of ChR2 expression and optrode placement (n=4 rats). **B**. Rasters and peristimulus time histograms of a representative single neuron showing increased firing rate during laser illumination (8–10mW, 473nm, 30s ON, 30s OFF, 4 Hz, 5 trials). **C**. *Left:* Proportion of neurons showing an increase (gold, n=43), decrease (blue, n=27), or no change (grey, n=42) in firing rate with laser ON. *Right:* Average firing rate at baseline (dark grey) and 4 Hz photoactivation for neurons showing increased (gold) changes in firing rate. **D**. Rasters and peristimulus time histograms of a representative single neuron showing increased firing rate during laser illumination (8–10mW, 473nm, 30s ON, 30s OFF, 2 Hz, 5 trials). **E**. *Left:* Proportion of neurons showing an increase (n=35), decrease (n=26), or no change (n=15) in firing rate with laser ON. *Right:* Average firing rate at baseline, and 2 Hz photoactivation for neurons showing increased changes in firing rate. All data are shown as mean ± SEM.

We next infused ChR2 into either the rPL or cPL and began avoidance conditioning 3–4 weeks after AAV infusion (Figure 6A). Following 10 days of avoidance training, rats were exposed to two tones presented in the absence of shock. PL neurons were illuminated throughout the first tone (4 Hz, 30 sec), followed by the second tone with the laser off. Photoactivation of rPL neurons at 4 Hz markedly reduced avoidance expression as reflected in the total time spent on the platform during the tone (Figure 6B; eYFP-control, n=9, 87% vs. eYFP-ChR2, n=14, 27%, unpaired t-test, t_(21)_=-4.779, p<0.001, Bonferroni corrected). In contrast to rPL, photoactivation of cPL had no effect on avoidance (Figure 6C&E).

**Figure 6.**
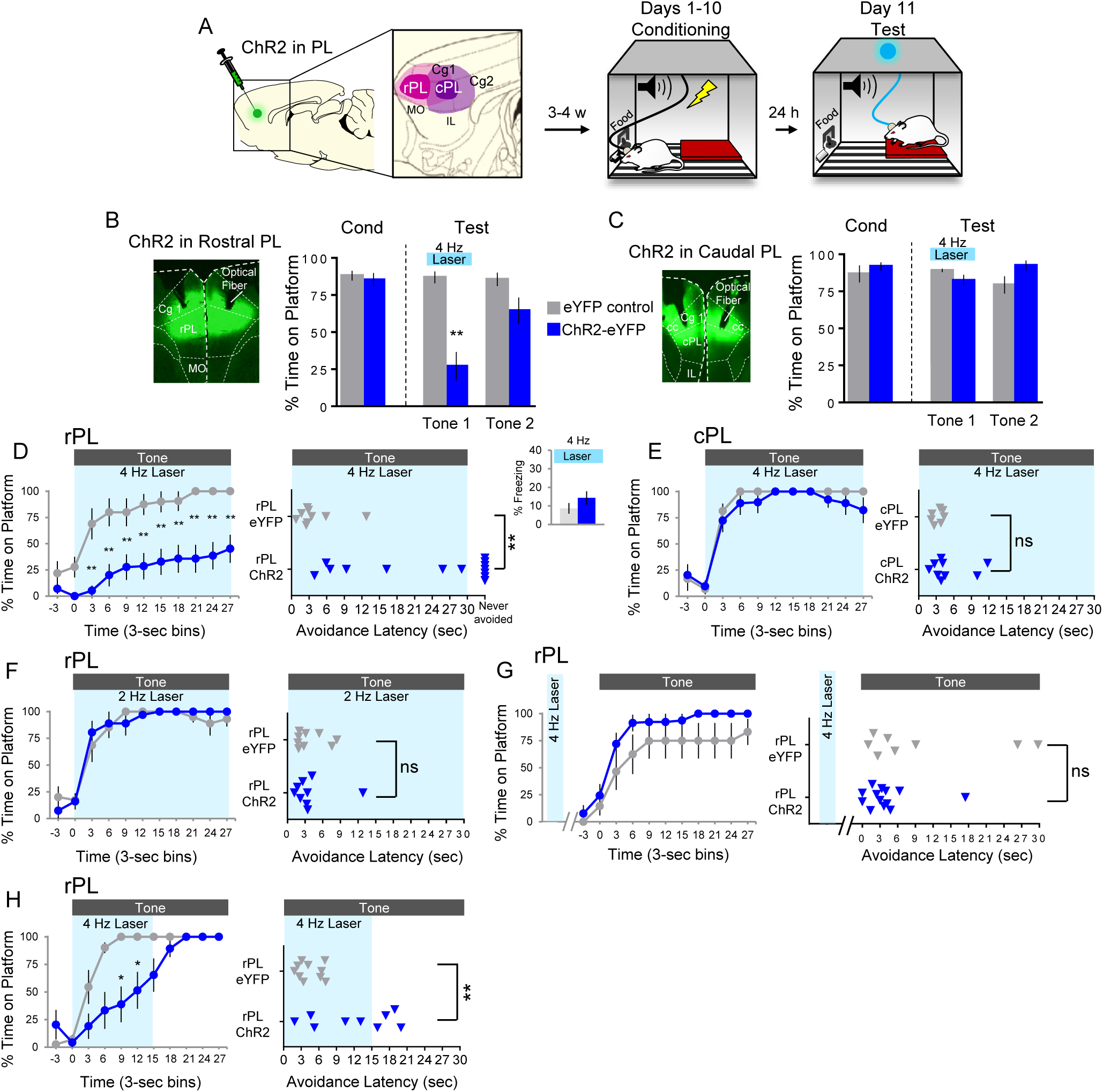
4 Hz photoactivation of rostral PL neurons during the tone delays or prevents avoidance. **A**. Schematic of viral infusion and location of min/max spread of AAV expression in rPL (pink) and cPL (purple), followed by avoidance training and test. At Test, 473nm light was delivered to rPL or cPL during the first 30-second tone presentation (Tone 1). **B**. *Left*: Micrograph of ChR2 expression and optical fiber placement in rPL. *Right:* Percent time on platform at Cond (Day 10, Tone 1) and Test (Day 11, Tone 1 with laser ON and Tone 2 with laser OFF) for eYFP-rPL control rats (grey, n=9) and ChR2-rPL rats (blue, n=14). **C**. *Left:* Micrograph of ChR2 expression and optical fiber placement in cPL. *Right:* Percent time on platform during Cond and Test for eYFP-cPL control rats (grey, n=7) and ChR2-cPL rats (blue, n=9). **D**. *Left:* Time spent on platform in 3 sec bins (Tone 1 at Test) revealed that rPL-ChR2 rats were significantly delayed in their avoidance compared to eYFP controls (repeated measures ANOVA, post hoc Tukey test). *Right:* Latency of avoidance for each rat (Mann Whitney U Test, Tone 1 at Test). 7/14 rats never avoided. *Inset:* 4 Hz photoactivation in rPL had no effect on freezing (Tone 1 at Test). **E**. Timeline of avoidance (left) and latency (right) for eYFP-cPL control ChR2-cPL rats revealed no effect of 4 Hz photoactivation of caudal PL. **F**. Timeline of avoidance (left) and latency (right) for eYFP-rPL control rats (grey, n=9) and ChR2-rPL rats (blue, n=9) revealed no effect of 2 Hz photoactivation. **G**. Timeline of avoidance (left) and latency (right) for eYFP-rPL control rats (grey, n=8) and ChR2-rPL rats (blue, n=13) revealed no effect of 4Hz photoactivation during a 30 sec ITI period. **H**. Timeline of avoidance (left) and latency (right) for eYFP-rPL control rats (grey, n=10) and ChR2-rPL rats (blue, n=9) revealed a delay in the initiation of avoidance with 4Hz photoactivation during the first 15 sec of the tone (Mann Whitney U test for time course and avoidance latency). All data are shown as mean ± SEM; *p<0.05; **p<0.01.

A closer examination of the time course of avoidance showed that photoactivation of rPL significantly reduced avoidance throughout the tone (Figure 6D left; repeated measures ANOVA, main effect (Group), F_(1)_=18.642, p<0.001, interaction effect (GroupxBins) F_(9)_=1.156, p=0.326, post hoc Tukey test, last 9 3-sec bins, all p’s<0.01). Photoactivation increased the latency to avoid in 7/14 rats, and blocked avoidance entirely in 7/14 rats (Figure 6D right; Mann Whitney U test, p<0.001), but it did not affect freezing (Figure 6D inset; eYFP-control 9% vs eYFP-ChR2 14% time freezing, unpaired t-test, p=0.275). Reducing the rate of photoactivation to 2Hz in rPL eliminated the effects on time course and latency of avoidance (Figure 6F). Furthermore, shifting the 30 sec of 4 Hz photoactivation to the inter-tone interval did not impair avoidance (Figure 6G). Thus, the photostimulation-induced impairment of avoidance showed specificity with respect to location, time, and frequency. Finally, reducing the duration of 4 Hz photoactivation from 30 sec to the first 15 sec of the tone delayed, but did not prevent, avoidance as indicated by time on platform (Figure 6H left; Mann Whitney U test, p’s<0.05 at 9–15 sec) and avoidance latency (Figure 6H right, unpaired t-test, t_(17)_=3.363, p=0.0037). Photoactivation of rPL at 4 Hz had no effect on locomotion, as indicated by distance traveled in an open field test during a 30 sec period (eYFP-control n=11, 2.71 m vs. eYFP-ChR2, n=15, 2.25 m, unpaired t-test, p=0.356). Nor did it affect anxiety levels, as both groups spent a similar amount of time in the center of the open field (eYFP-control = 2.6727 sec vs. eYFP-ChR2 = 2.6733 sec, unpaired t-test, p=0.999). These findings are consistent with the hypothesis that initiation of avoidance depends on tone-induced inhibition in rPL neurons.

## Discussion

We investigated the neural mechanisms of the initiation of active avoidance. Whereas pharmacological inactivation of PL delayed initiation of avoidance, optogenetic silencing did not. Single-unit recordings revealed that initiation of avoidance was correlated with inhibitory, rather than excitatory, tone responses of rostral PL neurons. Consistent with this, optogenetically activating rPL neurons to oppose tone-induced inhibition delayed or prevented the initiation of avoidance. These findings add to the growing body of evidence showing that inhibition within PL is key for conditioned behavior (Ehrlich et al., 2009, Ciocchi et al., 2010, Sotres-Bayon et al., 2012, Sparta et al., 2014) and highlight the importance of using in vivo recordings to guide optogenetic manipulations of behavior.

Our findings extend previous findings that PL activity is necessary for avoidance expression (Bravo-Rivera et al., 2014) by showing that PL processing contributes more to early, rather than late, avoidance. Our findings appear to be at odds with prior cFos studies, showing that the expression of active avoidance is correlated with increased activity in PL (Martinez et al., 2013, Bravo-Rivera et al., 2015). Importantly, the excitatory responses we observed in PL neurons were associated with either sensory correlates of the tone, or motor correlates of platform entry, compared to naïve controls. Thus, previously observed increases in cFos expression may represent sensory and/or motor features of avoidance-like behavior rather than avoidance itself, which was correlated only with inhibitory responses in PL. This resembles recent findings in the basolateral amygdala, where increased activity in some neurons were correlated with the cessation of movement, irrespective of motivation or valence (Amir et al., 2015, Pare and Quirk, 2017).

Previous work on PL has focused on its necessity for the expression of freezing during fear conditioning (Baeg et al., 2001, Vidal-Gonzalez et al., 2006, Burgos-Robles et al., 2009, Sierra-Mercado et al., 2011). However, in platform-mediated avoidance, freezing is either not reduced (Bravo-Rivera et al., 2014) or is increased (present study) by pharmacological inactivation of PL, suggesting that avoidance training alters the freezing circuit. Prior manipulations of prefrontal, amygdala, and striatal areas dissociated the expression of avoidance from the expression of freezing in this task (Bravo-Rivera et al., 2014, Rodriguez-Romaguera et al., 2016). Furthermore, we observed that stimulation of PL at 4 Hz impaired avoidance without increasing freezing. Thus, impaired avoidance is not a result of freezing-induced blockade of avoidance (Lazaro-Munoz et al., 2010).

Neurons in rPL project to the ventral striatum (VS; Sesack et al., 1989, Vertes, 2004), another region necessary for avoidance in this task (Bravo-Rivera et al., 2014) as well as other avoidance tasks (Darvas et al., 2011, Ramirez et al., 2015, Hormigo et al., 2016). Inputs to VS from rPL may be necessary for the initiation of platform-mediated avoidance via disinhibition of VS output, similar to appetitive behaviors where inputs to VS must be inhibited to trigger foraging for food (Rada et al., 1997, Saulskaya and Mikhailova, 2002, Do-Monte et al., 2017). The VS also receives input from the basolateral amygdala (BLA; McDonald, 1991, Wright et al., 1996, Groenewegen et al., 1999, Pitkänen et al., 2000), and this projection was recently implicated in the expression of shuttle avoidance (Ramirez et al., 2015). One possibility is that inputs from both rPL and BLA projecting to VS may be involved in avoidance, with rPL inputs initiating early avoidance, and BLA inputs initiating late avoidance. Thus, as the tone progresses and shock becomes more imminent, direct BLA inputs to VS are recruited. More work is needed to test this hypothesis.

We delayed avoidance initiation by photostimulating at 4 Hz, which was the average decrease in firing rate in neurons showing inhibitory responses to the tone. This rate of stimulation is much lower than the 20+ Hz used in previous behavioral studies employing channelrhodopsin (Liu et al., 2012, Felix-Ortiz and Tye, 2014, Marcinkiewcz et al., 2016, Villaruel et al., 2017, Warlow et al., 2017). Moreover, we have previously observed that photoactivation of infralimbic prefrontal cortex at rates ≥10 Hz was needed to reduce conditioned freezing (Do-Monte et al., 2015a). As 4 Hz is close to the average firing rate of mPFC putative projection neurons (Jung et al., 1998, Baeg et al., 2001, Burgos-Robles et al., 2009, Sotres-Bayon et al., 2012), the delay in avoidance was likely due to a reduction of neuronal inhibition rather than excitation above baseline. However, an important caveat is that CamKIIα-expressing neurons were activated indiscriminately, and were not limited to neurons showing inhibitory responses to the tone. Thus, in addition to reducing inhibitory responses in one population of cells, we likely induced excitation in another population of cells that do not normally express inhibitory tone responses. Both mechanisms would have the effect of increasing tone-induced activity at rPL targets, which could account for the more robust effect of photoactivation vs. muscimol inactivation on the initiation of avoidance.

There are multiple sources of input to rPL capable of driving inhibition during the tone. PL receives input from the BLA, vHPC, and OFC (Bacon et al., 1996, Vertes, 2006, Hoover and Vertes, 2007). BLA inputs to rPL are tone-responsive and robust, however, they appear to be excitatory (Orozco-Cabal et al., 2006, Little and Carter, 2012, Little and Carter, 2013), communicating auditory conditioned responses (Senn et al., 2014, Cheriyan et al., 2016, Burgos-Robles et al., 2017). Indeed, inactivating BLA reduces the CS-responsiveness of PL neurons (Laviolette et al., 2005, Sotres-Bayon et al., 2012). A more likely candidate for inhibitory input is the ventral hippocampus (vHPC), as inactivation of this area increases conditioned tone responses of PL neurons (Sotres-Bayon et al., 2012). Consistent with this, other studies have demonstrated that vHPC targets interneurons in the prefrontal cortex (Gabbott et al., 2002, Ishikawa and Nakamura, 2003, Tierney et al., 2004). Whereas vHPC activity is necessary to acquire fear (Maren and Holt, 2004, Sierra-Mercado et al., 2011, Chen et al., 2016), it is unknown whether vHPC activity is necessary for avoidance.

Excessive avoidance of stimuli that are not dangerous is clinically relevant for PTSD and other anxiety disorders. Rodent PL is thought to be homologous to the dorsal anterior cingulate (dACC) in humans (Bicks et al., 2015, Heilbronner et al., 2016). Although it is difficult to link unit recording findings with BOLD responses in fMRI studies, decreased activity in human dACC was correlated with active avoidance (Schlund et al., 2015), and avoidance has been correlated with functional connectivity between rostral dACC and striatum (Collins et al., 2014). In PTSD patients, avoidance symptoms correlate with excessive activity in rostral dACC (Marin et al., 2016). Together with our findings, this suggests that compromised inhibitory control of rostral dACC may predispose individuals to express avoidance when it competes with a high behavioral cost, and/or when it is not urgent.

## Methods

### Subjects

A total of 156 adult male Sprague Dawley rats (Harlan Laboratories, Indianapolis, IN) aged 3–5 months and weighing 320–420 g were housed and handled as previously described (Bravo-Rivera et al., 2014). Rats were maintained on a restricted diet (18 g/day) of standard laboratory rat chow to facilitate pressing a bar for food on a variable interval schedule of reinforcement (VI-30). All procedures were approved by the Institutional Animal Care and Use Committee of the University of Puerto Rico School of Medicine in compliance with the National Institutes of Health guidelines for the care and use of laboratory animals.

### Surgery

Rats were anesthetized with isofluorane inhalant gas (5%) first in an induction chamber then positioned in a stereotaxic frame (Kopf Instruments, Tujunga, CA). Isofluorane (2–3%) was delivered through a facemask for anesthesia maintenance. For pharmacological inactivations, rats were implanted with 26-gauge double guide cannulas (Plastics One, Roanoke, VA) in the prelimbic prefrontal cortex (PL; +3.0 mm AP; ±0.6 mm ML; −2.5 mm DV, 0° angle) to bregma). For optogenetic experiments, rats were bilaterally implanted with 22-gauge single guide cannulas (Plastics One, Roanoke, VA) in the prelimbic prefrontal cortex (PL; +2.6-2.8 mm AP; ±1.50 mm ML; −3.40 mm DV to bregma, 15°angle). An injector extending 2 mm beyond the tip of each cannula was used to infuse 0.5 μl of virus at a rate of 0.05 μl/min. The injector was kept inside the cannula for an additional 10 min to reduce back-flow. The injector was then removed and an optical fiber (0.22 NA, 200 nm core, constructed with products from Thorlabs, Newton, NJ) with 1 mm of projection beyond the tip of each cannula was inserted for PL illumination. The guide cannula and the optical fiber were cemented to the skull (C&B metabond, Parkell, Brentwood, NY; Ortho Acrylic, Bayamón, PR). For unit recording experiments, rats were implanted with a moveable array of 9 or 16 microwires (50 μm spacing, 3×3 or 2 × 8, Neuro Biological Laboratories, Denison, TX) targeting regions of PL along the rostral-caudal axis. After surgery, triple antibiotic was applied topically around the surgery incision, and an analgesic (Meloxicam, 1 mg/Kg) was injected subcutaneously. Rats were allowed a minimum of 7 days to recover from surgery prior to behavioral training.

### Behavior

Rats were initially trained to press a bar to receive food pellets on a variable interval reinforcement schedule (VI-30) inside standard operant chambers (Coulbourn Instruments, Whitehall, PA) located in sound-attenuating cubicles (MED Associates, St. Albans, VT). Bar-pressing was used to maintain a constant level of activity against which avoidance and freezing could reliably be measured. Rats were trained until they reached a criterion of ≥15 presses/min. Rats pressed for food throughout all phases of the experiment.

For platform-mediated avoidance, rats were trained as previously described (Bravo-Rivera et al., 2014). Briefly, rats were conditioned with a pure tone (30 s, 4 kHz, 75 dB) co-terminating with a scrambled shock delivered through the floor grids (2 s, 0.4 mA). The inter-trial interval was variable, averaging 3 min. An acrylic square platform (14.0 cm each side, 0.33 cm tall) located in the opposite corner of the sucrose pellet–delivering bar protected rats from the shock. The platform was fixed to the floor and was present during all stages of training (including bar-press training). Rats were conditioned for 10 days, with 9 tone-shock pairings per day with a VI-30 schedule maintained across all training and test sessions. The availability of food on the side opposite to the platform motivated rats to leave the platform during the inter-trial interval, facilitating trial-by-trial assessment of avoidance. Once rats learned platform-mediated avoidance, rats underwent a 2-tone expression test (2 tones with no shock).

### Drug infusions

The GABA-A agonist muscimol (fluorescent muscimol, BODIPY TMR-X conjugate, Sigma-Aldrich) was used to enhance GABA-A receptor activity, thereby inactivating target structures. Infusions were made 45 min before testing at a rate of 0.2 μl/min (0.11 nmol/ 0.2 μl/ per side), similar to our previous studies (Do-Monte et al., 2015b, Rodriguez-Romaguera et al., 2016).

### Viruses

The adeno-associated viruses (AAVs; serotype 5) were obtained from the University of North Carolina Vector Core (Chapel Hill, NC). Viral titers were 4 × 10^12^ particles/ml for channelrhodopsin (AAV5:CaMKIIα::hChR2(H134R)-eYFP) and archaerhodopsin (AAV5:CaMKIIα::eArchT3.0-eYFP) and 3 × 10^12^ particles/ml for control (AAV5:CaMKIIα::eYFP). Rats expressing eYFP in PL were used to control for any nonspecific effects of viral infection or laser heating. The CaMKIIα promoter was used to enable transgene expression favoring pyramidal neurons (Liu and Jones, 1996). Viruses were housed in a −80° C freezer until the day of infusion.

### Laser delivery

Rats expressing channelrhodopsin (ChR2) in PL were illuminated using a blue diode-pump solid state laser (DPSS, 473 nm, 2 or 4 Hz, 5 ms pulse width, 8–10 mW at the optical fiber tip; OptoEngine, Midvale, UT), similar to our previous study (Do-Monte et al., 2015a). Rats expressing archaerhodopsin (Arch) in PL were bilaterally illuminated using a DPSS green laser (532 nm, constant, 10–12 mW at the optical fiber tip; OptoEngine). For both ChR2 and Arch experiments, the laser was activated at tone onset and persisted throughout the 30 s tone presentation. Laser light was passed through a shutter/coupler (200 nm, Oz Optics, Ontario, Canada), patchcord (200 nm core, ThorLabs, Newton, NJ), rotary joint (200 nm core, 2 × 2, Doric Lenses, Quebec city, Canada), dual patchcord (0.22 NA, 200 nm core, ThorLabs), and bilateral optical fibers (made in-house with materials from ThorLabs and Precision Fiber Products, Milpitas, CA) targeting the specific subregions in PL. Rats were familiarized with the patchcord during bar press training and during the last 4 d of avoidance training before the expression test.

### Single-unit recordings

Rats implanted with moveable electrode arrays targeting PL were either avoidance conditioned as previously described or exposed to the training environment (platform, tone presentations, behavior box) in the absence of the shock. Extracellular waveforms that exceeded a voltage threshold were digitized at 40 kHz and stored on a computer. Waveforms were then sorted offline using three-dimensional plots of principal component and voltage vectors (Offline Sorter; Plexon, Dallas, TX) and clusters formed by individual neurons were tracked. Timestamps of neural spiking and flags for the occurrence of tones and shocks were imported to NeuroExplorer for analysis (NEX Technologies, Madison, AL). Because we used a high impedance electrode in the current study (~750–1000 kOhm), we were unable to sample interneurons. Data was recorded during the entire session except during the 2 sec shock. After conditioning, rats were tested for avoidance expression. For avoidance assessment, rats received full conditioning sessions (with shocks) across days. Inclusion of the shock prevented extinction of avoidance. After each day, electrodes were lowered 150 ^M to isolate new neurons for the following session the next day. To detect tone-elicited changes in PL activity, we assessed whether neurons changed their firing rate significantly during the first 500–1000 ms after tone onset across the first 5 trials. A Z-score for each 500 ms bin was calculated relative to 20 pre-tone bins of equal duration (10 sec pre-tone). PL neurons were classified as showing excitatory tone responses if the initial bins exceeded 2.58 z’s (p < 0.01, two-tailed). PL neurons were classified as showing inhibitory tone responses across time if any of the initial two tone bins exceeded −1.96 Z’s (p < 0.05, two-tailed). To detect changes in PL activity during platform entry, we employed the same procedure used for assessing tone responses. We assessed whether neurons changed their firing rate significantly during the first 500–1000 ms after platform entry. A Z-score for each 500 ms bin was calculated relative to the same pre-tone baseline. Heat maps of single unit data were generated with Z-scores from baseline through the 28 sec after tone onset or platform entry.

### Optrode recordings

Rats expressing Arch or ChR2 in PL were anesthetized with urethane (1g/Kg, i.p.; Sigma Aldrich) and mounted in a stereotaxic frame. An optrode consisting of an optical fiber surrounded by 8 or 16 single-unit recording wires (Neuro Biological Laboratories) was inserted and aimed at PL (AP, +2.8 mm; ML: −0.5; DV: −3.5). The optrode was ventrally advanced in steps of 0.03 mm. Single-units were monitored in real time (RASPUTIN, Plexon). After isolating a single-unit, a 532 nm laser was activated for 10 sec within a 20 sec period, at least 10 times for Arch-infected PL neurons. For ChR2-infected PL neurons, a 473 nm laser was activated for 30s at a rate of 2 or 4 Hz (5 ms pulse width) within a 90s period (60s ITI), at least 5 times. Single-units were recorded and stored for spike sorting (Offline Sorter, Plexon) and spike-train analysis (Neuorexplorer, NEX Technologies). Excitatory and inhibitory responses were calculated by comparing the average firing rate of each neuron during the 10 sec of laser OFF with the 10 sec of laser ON for Arch neurons and during 30 sec laser OFF just prior to the 30 sec of laser ON for ChR2 neurons (Paired t-test, 1 s bins).

### Open field task

Locomotor activity in the open field arena (90 cm diameter) was automatically assessed (ANY-Maze) by comparing the total distance travelled between 30 sec trials (laser off versus laser on), following a 3 min acclimation period for optogenetic experiments. The distance traveled was used to assess locomotion and time in center was used to assess anxiety. For pharmacological inactivation experiments, distance traveled and time in center was measured over a 3 min period following a 3 min acclimation period 45 min after MUS or SAL was infused prior to sacrificing animals.

### Histology

After behavioral experiments, rats were deeply anesthetized with sodium pentobarbital (450 mg/kg i.p.) and transcardially perfused with 0.9 % saline followed by a 10 % formalin solution. Brains were removed from the skull and stored in 30 % sucrose for cryoprotection for at least 72 h before sectioning and Nissl staining. Histology was analyzed for placement of cannulas, virus expression, and electrodes.

### Data Collection and Analysis

Behavior was recorded with digital video cameras (Micro Video Products, Peterborough, Ontario, Canada). Freezing and platform avoidance was quantified by observers blind to the experimental group. Freezing was defined as the absence of all movement except for respiration. Avoidance was defined as the rat having at least three paws on the platform. In a subset of animals, AnyMaze software was available for recording and calculating freezing and avoidance (Stoelting, Wood Dale, IL). The time spent avoiding during the tone (percent time on platform) was used as our avoidance measure. Avoidance and freezing to the tone was expressed as a percentage of the 30 second tone presentation. Our experimental groups typically consisted of approximately 15 animals. This is typical of other laboratories and results in sufficient statistical confidence. Moreover, it also agrees with the theoretical minimum sample size given by:

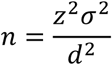

where z = the level of confidence desired (in standard deviations), σ = the estimate of the population standard deviation, and d = the acceptable width of the confidence interval. Technical replications, testing the same measurement multiple times, and biological replications, performing the same test on multiple samples (individual rats or single units), were used to test the variability in each experiment. Statistical significance was determined with Student’s two-tailed t-tests, Fisher Exact tests, Chi Square tests, Mann Whitney U tests, or repeated-measures ANOVA, followed by post hoc Tukey analysis, and Bonferroni corrections, where appropriate using STATISTICA (Statsoft, Tulsa, OK) and Prism (Graphpad, La Jolla, CA).

## Acknowledgements

This study was supported by NIH grants F32-MH105185 to MMD, R36-MH102968 to CBR, R36-MH105039 to JRR, R25-NS080687 to PAPR, R37-MH058883 and P50-MH106435 to GJQ, and the University of Puerto Rico President’s Office. We thank Drs. Denis Pare and Drew Headley for comments on an earlier version. We also thank Jorge Maldonado de Jesus, Valeria Lozada-Miranda, Joyce Mendoza-Navarro, Jorge Iravedra-Garcia, and Fabiola Gonzalez-Díaz for help with behavioral experiments and Carlos Rodríguez and Zarkalys Quintero for technical assistance.

---

**Video 1.4 Hz photoactivation of rostral PL neurons during the tone impairs avoidance.**

Video of an individual rat with ChR2 infused into rPL showing avoidance behavior on the last day of avoidance training (Day 10) at Tone 1, followed by the rat’s behavior at Test (Day 11) with the laser on during the tone (4 Hz, 30 sec duration, 5 ms pulse width, 8–10mW light intensity).

